# Targeting conserved sequences circumvents the evolution of resistance in a viral gene drive against human cytomegalovirus

**DOI:** 10.1101/2021.01.08.425902

**Authors:** Marius Walter, Rosalba Perrone, Eric Verdin

## Abstract

Gene drives are genetic systems designed to efficiently spread a modification through a population. They have been designed almost exclusively in eukaryotic species, and especially in insects. We recently developed a CRISPR-based gene drive system in herpesviruses that relies on similar mechanisms and could efficiently spread into a population of wildtype viruses. A common consequence of gene drives in insects is the appearance and selection of drive-resistant sequences that are no longer recognized by CRISPR-Cas9. Here, we analyze in cell culture experiments the evolution of resistance in a viral gene drive against human cytomegalovirus. We report that after an initial invasion of the wildtype population, a drive-resistant population is positively selected over time and outcompetes gene drive viruses. However, we show that targeting evolutionary conserved sequences ensures that drive-resistant viruses acquire long-lasting mutations and are durably attenuated. As a consequence, and even though engineered viruses don’t stably persist in the viral population, remaining viruses have a replication defect, leading to a long-term reduction of viral levels. This marks an important step toward developing effective gene drives in herpesviruses, especially for therapeutic applications.

## Introduction

Herpesviruses are nuclear-replicating viruses with large dsDNA genomes (100-200 kb) that encode hundreds of genes (1). They establish life-long persistent infections and are implicated directly or indirectly in numerous human diseases (2). In particular, human cytomegalovirus (hCMV) is an important threat to immunocompromised patients, such as HIV-infected individuals, receivers of organ transplants and unborn infants.

The use of defective viruses that interfere with the replication of their infectious parent after co-infecting the same cells – a therapeutic strategy known as viral interference – has generated a lot of interest in recent years. Different strategies against HIV, influenza or zika viruses have been described (3–5) The viral gene drive system that we recently reported in herpesviruses (with hCMV) represents a novel interfering strategy that causes the conversion of wildtype viruses into new recombinant viruses and drives the native viral population to extinction (6). Gene drives are genetic modifications designed to spread efficiently in a target population (7–10). They have been engineered principally in insect species, such as mosquitoes, and are seen as a potential strategy to eradicate vector-borne diseases, such as malaria and dengue. Most gene drives rely on CRISPR-mediated homologous recombination and have been restricted to sexually reproducing organisms. The viral gene drive system that we developed with hCMV relies on co-infection of cells by a wildtype and an engineered virus (6). Cleavage by Cas9 and repair by homologous recombination lead to the conversion of wildtype viruses into new recombinant viruses (Figure 1A). We demonstrated that gene drive viruses can replace their wildtype counterparts and spread in the viral population in cell culture experiments. Moreover, we showed that a gene drive virus presenting severe replicative defects could spread into the viral population and ultimately reduce viral levels. This development represents a novel therapeutic strategy against herpesviruses.

**Figure 1:**
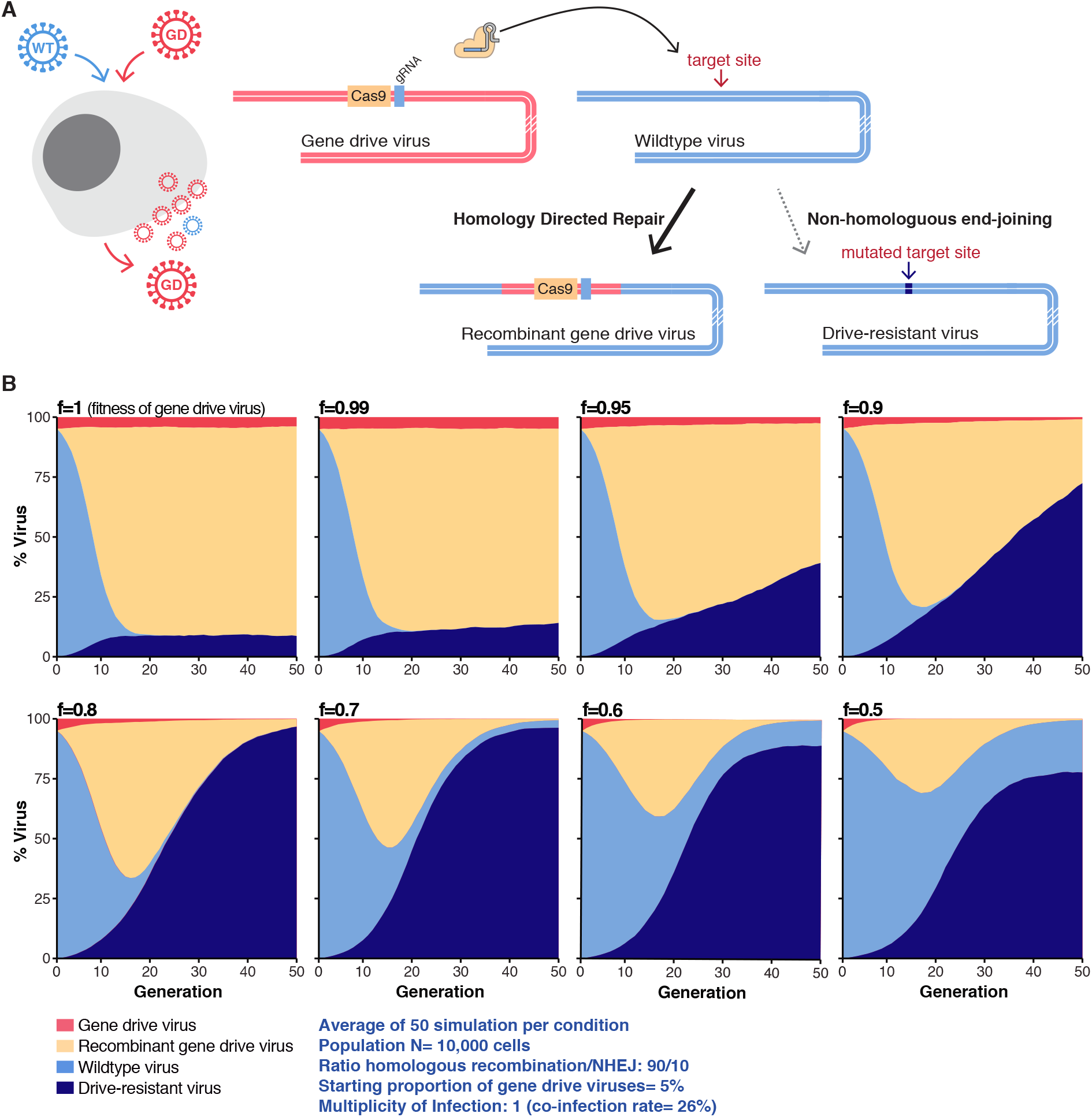
Evolution of gene drive resistance in numerical simulations. **A.** CRISPR-based gene drive sequences are, at a minimum, composed of *Cas9* and a gRNA targeting the complementary wildtype locus. After co-infection of a cell by a wildtype and a gene drive virus, Cas9 targets and cleaves the wildtype sequence. Homology-directed repair of the damaged wildtype locus using the gene drive sequence as a repair template causes the conversion of the wildtype locus into a new gene drive sequence and the formation of new recombinant viruses. In 5–10% of cases, repair of the damaged genome by non-homologous end-joining (NHEJ) creates drive-resistant viruses with a mutated target site. **B**. Numerical simulation showing that the appearance and positive selection of drive-resistant viruses depend on the replicative fitness f of gene-drive viruses. In these simulations, at each viral generation, N virtual cells were randomly infected and co-infected by N viruses, producing a new generation of viruses. When a cell is co-infected by wildtype and gene-drive viruses, wildtype viruses are converted to new gene drive viruses or resistant viruses in a 90/10 ratio.

An important challenge lies in designing gene drive viruses with replicative defects that still spread efficiently in the viral population and ultimately reduce viral levels. In the ideal scenario, gene drive viruses are non-infectious but are complemented by wildtype factors upon co-infection and can spread in the population only as long as wildtype viruses are present. Our initial study targeted *UL23*, a viral gene that encodes a tegument protein involved in immune evasion. UL23 is dispensable in normal cell culture conditions, but necessary in presence of interferon γ (6, 11, 12). The tegument of herpesvirus is a layer of protein that lies between the genome-containing capsid and the viral envelope (1, 13). Tegument proteins are released in the cell upon viral entry and are often involved in transcriptional activation and immune evasion. Tegument proteins are attractive targets for a viral gene drive because many of them function early in the viral cycle – where they could be complemented by a co-infecting wildtype virus – but only accumulate during later stages of the infection. In the present study, we designed gene drives against several tegument genes and showed that they indeed represent effective targets.

An important aspect of gene drive in sexually reproducing organisms, such as mosquitos, is the appearance and selection of drive-resistant alleles that are no longer recognized by the CRISPR guide RNA (gRNA) (14–16). Such alleles can already exist in the wild population or appear when the target site is repaired and mutated by non-homologous end-joining instead of homologous recombination (Figure 1A). These sequences are “immune” to future conversion into the drive allele, are often positively selected over time, and limit the ability to permanently modify a wildtype population (16–19).

In this report, we sought to investigate the appearance and evolution of resistance in a viral gene drive. Through numerical simulations and cell culture experiments with hCMV, we show that, after the initial propagation, a drive-resistant viral population is positively selected and outcompetes gene drive viruses over time. By designing and testing multiple gene drives that disable critical viral genes, we show that targeting evolutionary conserved sequences ensures that drive-resistant viruses have a replication defect that leads to a longterm reduction of viral levels.

## Results

### Drive-resistant viruses are positively selected over time

We first attempted to model the evolution of gene drive resistance using numerical simulations. Our initial study indicated that after the successful propagation of the gene drive in the wildtype population, around 5-10% of viruses had accumulated mutations of the target site. These mutations were caused by non-homologous repair of the cleavage site and rendered the mutated viruses resistant to the drive (6). Using a 10% estimate and considering in this scenario that resistant viruses would replicate at the same level as wildtype, we modeled the evolution of the viral population over time (Figure 1B, Supplementary Figure S1). These simulations showed that, depending on the replicative fitness of gene drive viruses, a population of drive-resistant viruses is often positively selected over time. At low fitness cost, the gene drive virus first spread efficiently in the wildtype population, but is outcompeted in the long run. At high fitness cost, a resistant population appears quickly and gene drive viruses disappear rapidly. Overall, these simulations predict that the appearance of drive-resistant viruses would prevent the gene drive from permanently remaining in the viral population.

We next evaluated experimentally the evolution of resistance in cell culture. Our initial experimental system involved an unmodified hCMV virus expressing GFP (strain Towne, referred hereafter as Towne-GFP) and an mCherry-expressing gene drive virus (GD-UL23) targeting *UL23* (6) (Figure 2A). *UL23* is dispensable in normal cell culture conditions (11, 12), but the GD-UL23 virus was built in a different viral strain (TB40/E) and replicated significantly slower than Towne-GFP (p=0.0011, Welsh t-test on log-transformed data. Figure 2B). Human foreskin fibroblasts were co-infected at a low multiplicity of infection (MOI=0.1) with the two viruses in different starting proportions, and the infection was subsequently propagated for several weeks by infecting fresh cells every 10-15 days. The MOI was kept around 0.1 at each passage. The proportion of the different viruses was evaluated over time by plaque assay (Figures 2C-D). The mCherry reporter enabled us to follow the spread of the gene drive in the viral population: viruses expressing mCherry represent gene drive viruses, and viruses expressing only GFP are unconverted – either unmodified or drive-resistant – viruses. In each of 12 biological replicates and independently of the starting proportion of GD-UL23, wildtype viruses were first converted to new gene drive viruses, and the population of GFP-only viruses reached a minimum level of 5–40% after 10–40 days (Figures 2C). In a second phase, however, GFP-only viruses appeared to be positively selected, and their proportion rebounded until they represented the majority of the viral population. This showed that in the long term, drive-resistant viruses are positively selected.

**Figure 2:**
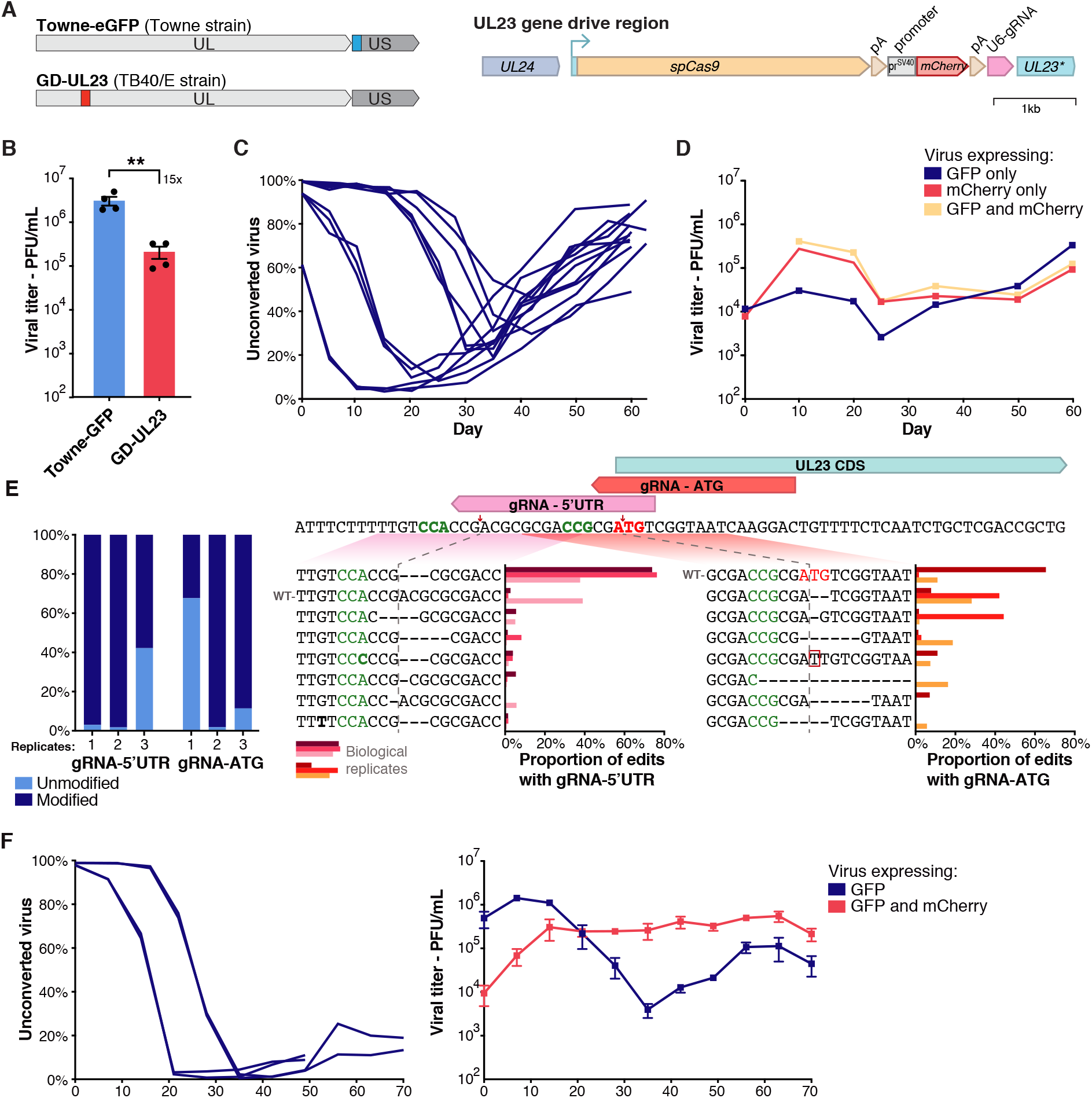
Evolution of resistance in a gene drive against UL23. **A.** Left: Localizations of gene drive and GFP cassettes on hCMV genomes. UL/US indicates Unique Long/Short genome segments. Right: the gene drive cassette comprises *spCas9* under the control of *UL23* endogenous promoter, followed by an SV40 polyA signal, an SV40 promoter driving an mCherry reporter, a beta-globin polyA signal and a U6-driven gRNA. **B**. Viral titer in fibroblasts infected with Towne-GFP or GD-UL23 viruses after 7 days, as measured by plaque assay. MOI=1, n=4. **: p < 0.01, Welsh t-test on log-transformed data. **C**. Proportion over time of viruses expressing only GFP after co-infection with Towne-GFP and GD-UL23, as measured by plaque assay (MOI=0.1 at day 0). Every line corresponds to a biological replicate (n=12), with different starting proportions of GD-UL23: 40%, 10%, 1% or 0.1%, 3 replicates per condition. **D.** Viral titers over time of viruses expressing GFP alone, mCherry alone, or both, after co-infection with Towne-GFP and GD-UL23. This panel corresponds to the starting proportion of 40% that is shown in C. n=3, **E**. Amplicon sequencing of the CRISPR target site at the end of the drive (corresponding to experiments presented in supplementary figure 2). Left: proportion of edited genomes, right: Relative contribution of each edit. CRISPR cleavage site is indicated by a red arrow, the protospacer adjacent motif (PAM) is highlighted in green, and *UL23* start codon in red. n=3. **F**. Proportion over time of viruses expressing only GFP (left) and viral titers (right) after co-infection with Towne-GFP and GD^Towne^-UL23. MOI=1 at day 0, n=4. Titers are expressed in PFU (plaque-forming unit) per mL of supernatant. Data shows mean and SEM between biological replicates.

These first experiments were carried out with two viruses from different strains, which complicate the interpretation of the results. To alleviate the influence of the viral strain, we analyzed the evolution of resistance in a strict Towne background. Two gene drive plasmids (with a gRNA targeting either *UL23* 5’UTR or *UL23* start codon) were separately transfected into fibroblasts, and cells were subsequently infected with Towne-eGFP. The population of mCherry-expressing recombinant viruses and of GFP-only (unmodified or resistant) viruses was followed over time. As observed previously, the gene drive cassette spread efficiently until the population of GFP-only viruses reached a minimum after 70 days (Supplementary Figure S2A). Introduction of a cassette in the viral genome by transfection of a plasmid is a notoriously inefficient process. It was nonetheless sufficient to generate enough recombinant gene drive viruses to enable propagation of the gene drives through the viral population, which further demonstrated the high efficiency of the system. Importantly, directly transfecting a plasmid circumvented the time-consuming process of generating and purifying recombinant viruses, allowing to test multiple gene drive constructs in parallel. The population of drive-resistant viruses at 70 days was characterized by amplicon sequencing of the target site, in three biological replicates for the two conditions (Figure 2E, Supplementary Figure S2B). As expected, the target site was heavily mutated, with up to 98% of sequences modified in some of the replicates. Interestingly, around 80% of sequences edited with gRNA-5’UTR had the same 3pb deletion. By contrast, edits generated by gRNA-ATG were more diverse, with a dominant 2pb deletion. These results confirmed that drive-resistant viruses accumulate over time, limiting the impact of the drive.

Finally, we isolated and purified a gene drive virus against *UL23* in a Towne strain (GD^Towne^-UL23). Fibroblasts were co-infected with Towne-GFP and GD^Towne^-UL23 (MOI=1), and the population of unconverted viruses expressing only GFP was analyzed over time. Cells were infected with a starting proportion of GD^Towne^-UL23 of 1-2%, and the infection was propagated for several weeks by infecting fresh cells every 7-8 days (MOI around 1 at each passage). In this experiment, the drive achieved more than 99% penetrance in the viral population, and the proportion of unconverted viruses only rebounded slightly and plateaued at around 10–20% (Figure 2F, left panel). UL23 is dispensable in normal cell culture conditions (6, 11), and viral titers remained constant over the course of the experiment (Figure 2F, right panel). This result showed that a gene drive without fitness cost can stably invade the wildtype population, as predicted by numerical simulations.

These numerical and experimental results indicate that, after the initial invasion of the wildtype population, a drive-resistant viral population is positively selected over time and outcompetes gene drive viruses. Importantly, however, we showed that, in the absence of associated fitness costs, gene drive sequences can reach almost full penetrance and be maintained durably in the viral population.

### Gene drives against conserved regions of *UL26* and *UL35* spread efficiently in the viral population

We next sought to design gene drives that would lead to a long-term reduction of viral levels. Drive-resistant viruses are created by imperfect repair of the CRISPR cleavage site. We reasoned that the appearance and selection of drive-resistant viruses could be circumvented if the mutation rendered viruses nonfunctional, for example if it knocked-out a critical viral gene. In this case, numerical simulations predicted that drive-resistant viruses would also be counter-selected and would not accumulate (Supplementary Figure S3), leading to a longterm reduction of viral titers. We, therefore, designed several gene drives targeting hCMV genes that are necessary for efficient viral replication (summary in Table 1). The gene drive cassette was inserted in the coding sequence of the viral gene. In addition, CRISPR gRNAs were designed in evolutionary conserved sequences, so that any mutation would potentially affect viral fitness. In this situation, both gene drive and drive-resistant viruses would have a replication defect, leading to a long-term reduction of viral levels.

**Table 1:** Summary of gene drive experiments against several viral genes

| Gene | Function | Phenotype | Ability to drive |
| --- | --- | --- | --- |
| *UL23* | Tegument gene, involved in evasion of the innate immune response (11) | Dispensable (11) | Drive efficiently, up to 99% of conversion |
| *UL122* | Encode Immediate early protein IE2, responsible for the initiation of viral replication (20) | Essential (20) | Do not drive, no recombinant viruses could be observed |
| *UL79* | Regulation of transcription of late viral transcript (28) | Essential (21) | Do not drive, very few recombinant viruses |
| *UL99* | Tegument gene, essential for viral envelopment (22) | Essential (22) | Do not drive, no recombinant virus |
| *UL55* | Encode fusion protein gB | Essential (23) | Do not drive, no recombinant virus |
| *UL26* | Tegument gene, involved in immune evasion, transcriptional activation and virion stability (24, 29–31) | Severe growth defect (24) | Drive efficiently |
| *UL35* | Tegument gene, important for viral replication and virion formation (32) | Moderate growth defect (26) | Drive efficiently |
| *UL69* | Tegument gene, involved in unspliced mRNA nuclear export and cellular arrest (25, 33) | Severe growth defect (25) | Very limited drive at high MOI |
| *UL82* | Tegument gene, stimulate immediate-early transcription and inhibit host innate response (27, 34) | Moderate growth defect (27) | Very limited drive at high MOI |

We first attempted to build gene drive viruses against *UL122* (IE2), *UL79, UL99* and *UL55* (gB), using fibroblasts stably expressing the different viral genes. These viral genes are essential for viral replication and mutant viruses are non-infectious (20–23). We could generate gene drive viruses against *UL79* and *UL99* in complementing cells, but we observed only very few second-generation recombinant viruses when co-infecting with unmodified Towne-GFP. We were not able to build gene drive viruses against *IE2* and gB, in part because we didn’t succeed to generate fibroblasts stably expressing these two proteins. We also transfected fibroblasts with each individual plasmid before infecting cells with Towne-GFP virus, but we once again didn’t observe any recombinant viruses. These first attempts were unsuccessful, which suggested that co-infection with Towne-GFP was not sufficient to rescue these very strong loss-of-function mutants (Table 1). We next sought to target viral genes that are not absolutely essential, but still required for efficient viral replication. As explained in the introduction, we hypothesized that tegument proteins represented attractive targets. We designed gene drive plasmids against the tegument genes *UL26, UL35, UL69* and *UL82*, which, when mutated or deleted, lead to moderate growth defects (24–27). Fibroblasts were independently transfected with each individual plasmid, infected with Towne-GFP virus (MOI=1) and the population of recombinant viruses was followed over time (Table 1). We observed that the constructs against *UL26* and *UL35* could spread in the wildtype population, and these genes were selected for subsequent experiments.

CRISPR gRNAs against *UL26* and *UL35* were chosen in sequences evolutionarily conserved at the DNA and amino-acid levels. UL26 is a tegument protein involved in immune evasion, transcriptional activation and virion stability (24, 29–31). It is a member of the US22 family of herpesviral proteins (35), a family of proteins with conserved motifs across several herpesviruses. We aligned 31 protein sequences of US22 family members and screened for conserved motifs (Figure 3A, Supplementary Figure S4). Two highly conserved motifs were found, and we chose to design a gRNA that would disrupt a very conserved proline in the second motif. In parallel, available DNA sequences for 235 hCMV clinical and laboratory viruses were aligned, and the frequency of variants compared to the Towne reference was calculated (Figure 3B). The gRNA against *UL26* was chosen in a region of low variation, with less than 1% of sequenced hCMV viruses having a polymorphism in the first 18 bp of the gRNA sequence. A highly variable site was found in the gRNA sequence, but at a position (the 19th base) that presumably doesn’t efficiently prevent DNA cleavage (36). Most repairs of the predicted cut site would disrupt the evolutionary conserved proline. The gene drive cassette comprised *Cas9*, an mCherry reporter and a U6-driven gRNA (Figures 3C-D). Because the CRISPR target site against *UL26* was located far from the start codon (around 200 bp), a polyadenylation signal and a promoter with late kinetics (from hCMV *UL99*) were added to the gene drive cassette upstream of *Cas9*. The gRNA against *UL35* was designed in a similar manner. *UL35* encodes two isoforms with a common C-terminal domain but different N-termini (26, 37). Mutant viruses lacking the longer isoform have a modest replicative defect, and the gRNA against *UL35* targeted this first start codon, so that any mutations at the predicted cut site would abrogate translation of the long isoform (Figures 3D-F). This gRNA could not be designed in a region of low genetic variation (Figure 3E). This finding has little consequence for cell culture experiments, but suggests that this particular gene drive virus could be less efficient at targeting clinical hCMV strains.

**Figure 3:**
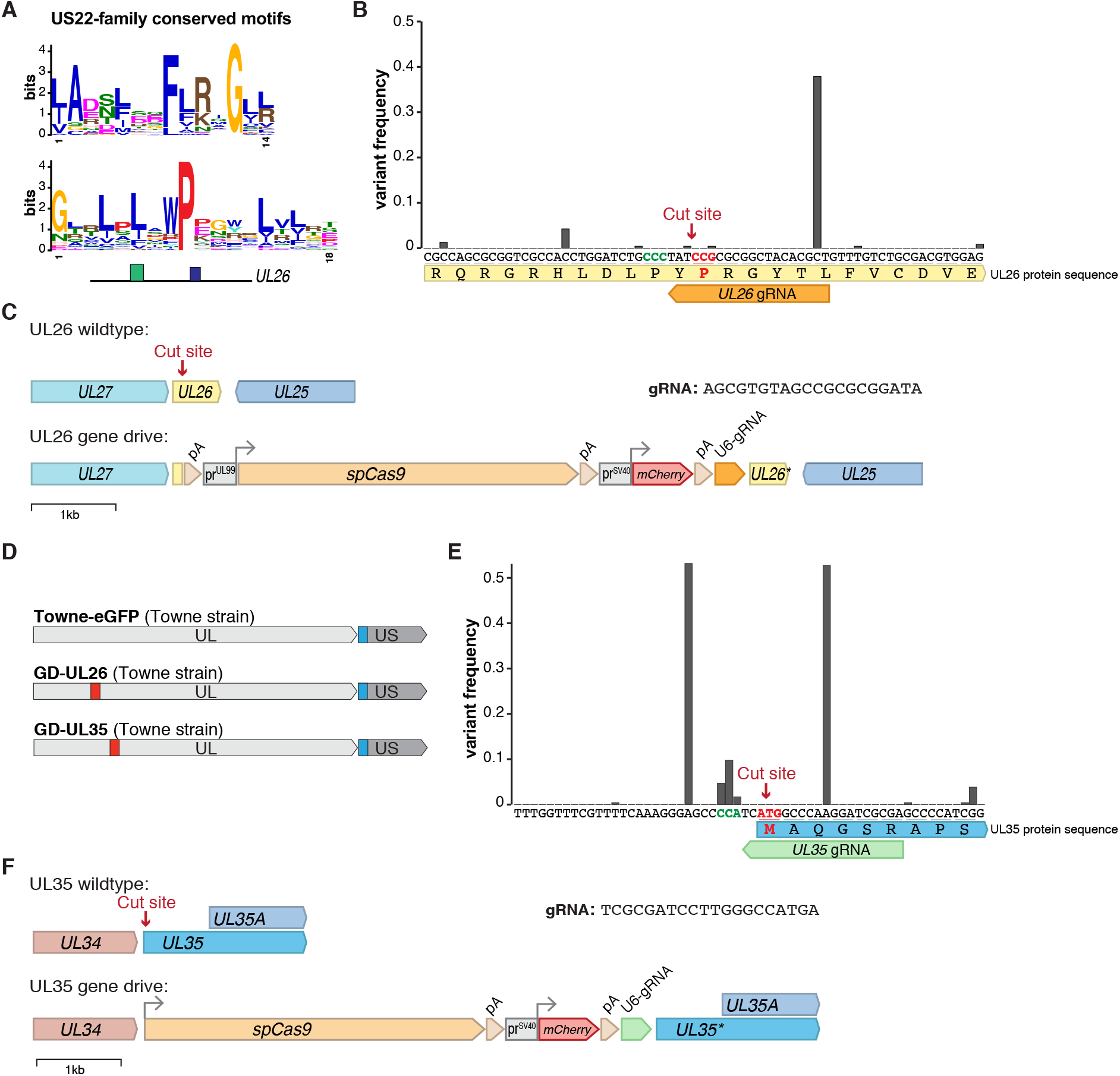
Design of gene drives against *UL26* and *UL35*. **A**. Logo sequences of the two most conserved motifs in the US22 family of herpesviral proteins. Bottom: localization of the two motifs on UL26 protein (blue: first motif, green: second motif). **B**. Frequency of genetic variants around *UL26* target site, from 235 hCMV strains. CRISPR protospacer adjacent motif (PAM) is highlighted in green, and the targeted proline codon in red. **C**. Wildtype and gene drive *UL26* regions. The gene drive cassette comprises an HSV1-TK polyA signal, *spCas9* under the control of an *UL99* late viral promoter, followed by a SV40 polyA signal, an SV40 promoter driving a mCherry reporter, a beta-globin polyA signal and a U6-driven gRNA. **D**. Localizations of gene drive and GFP cassettes on hCMV genomes. UL/US indicates Unique Long/Short genome segments. **E**. Frequency of genetic variants around UL35 target site, from 235 hCMV strains. CRISPR PAM is highlighted in green, and the targeted start codon in red. **F**. Wildtype and gene drive *UL35* region. The gene drive cassette is the same as in C, except that *spCas9* expression is controlled by *UL35* endogenous promoter.

Infectious gene drive viruses against *UL26* and *UL35* (GD-UL26 and GD-UL35, in Towne strain) were isolated and purified in fibroblasts. Of note, mutants lacking UL26 have severe growth defects (24) and the isolation and plaque assay of GD-UL26 viruses was carried on in cells stably expressing UL26. In wildtype fibroblast, GD-UL26 replication was almost completely abrogated compared to Towne-GFP while GD-UL35 replicated with a more moderate but significant ninefold growth defect (p<0.0001 and p=0.0039, respectively, ANOVA on log-transformed data. Figure 4A). GD-UL35 viral plaques were also significantly smaller compared to Towne-GFP, with a median plaque size 30% smaller (p=0.0003, Mann-Whitney test. Figure 4B).

**Figure 4:**
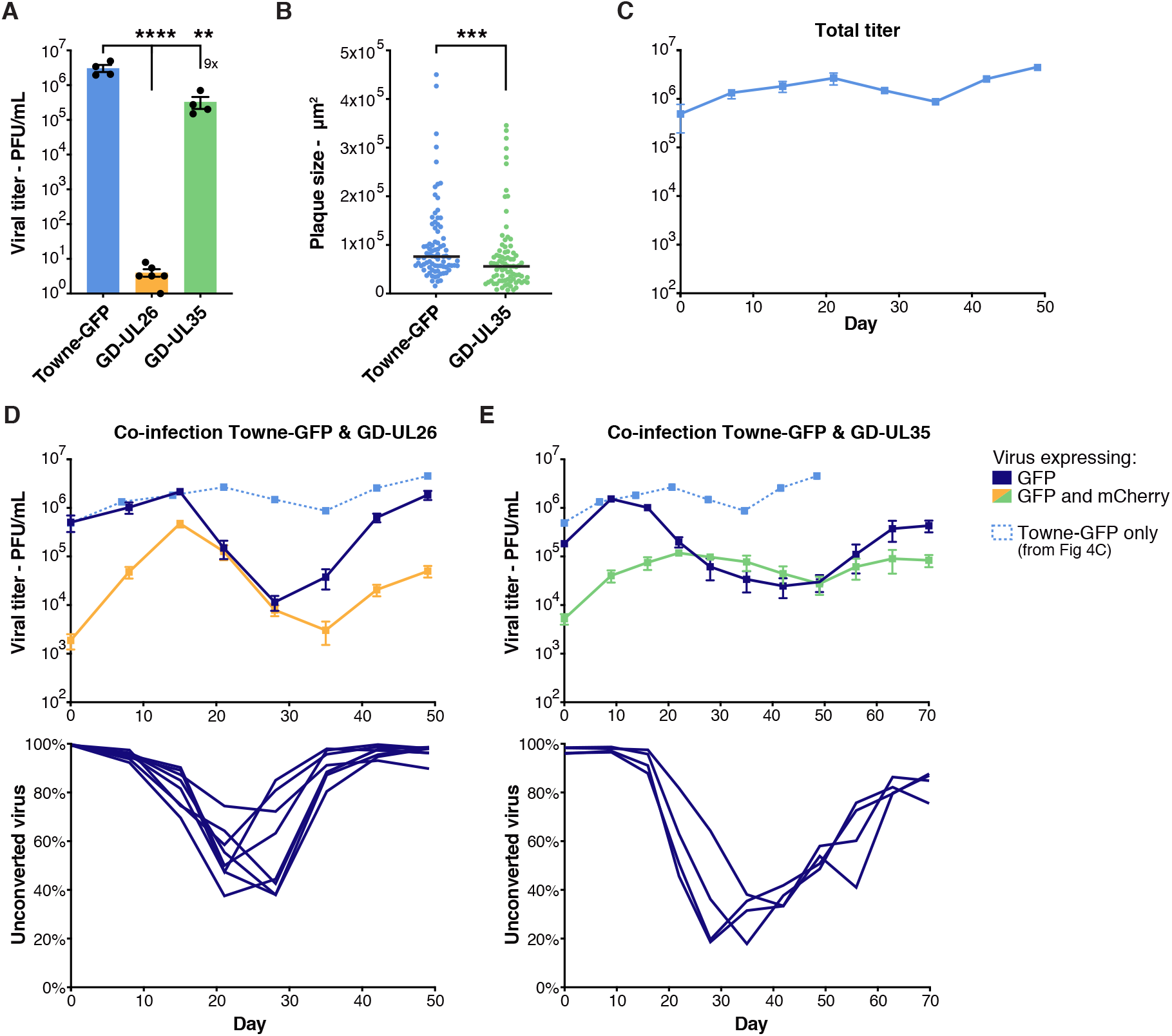
Gene drives against *UL26* and *UL35* spread in the viral population. **A.** Viral titer in fibroblasts infected with Towne-GFP, GD-UL26 or GD-UL35 viruses after 7 days. MOI=1, n=4. **B**. Plaque size in fibroblasts infected with Towne-GFP or GD-UL35, after 8 days. Black lines indicate the median, n=80. **C**. Viral titer over time in fibroblasts infected with Towne-GFP At each time point, supernatant was used to infect fresh cells to propagate the infection. n=4. **D-E**. Upper panels: Viral titer over time in fibroblasts co-infected with Towne-GFP and GD-UL26 (left) or GD-UL35 (right). Viruses either express GFP only (dark blue), or both GFP and mCherry (yellow/green).Viruses expressing mCherry represent gene drive viruses. Results from panel C showing mono-infection with Towne-GFP are superposed (light blue). Lower panels: proportion of the viral population expressing only GFP, representing unconverted (unmodified or drive-resistant) viruses. n=8 (GD-UL26) or n=4 (GD-UL35). Titers were measured by plaque assay and are expressed in PFU (plaque-forming unit) per mL of supernatant. Data shows mean and SEM between biological replicates. Asterisks summarize the results of statistical tests. **: p < 0.01, ***: p < 0.001, ****: p < 0.001. (A: one-way ANOVA with Dunnet’s multiple comparison test on log-transformed data, B: Mann-Whitney test.)

To analyze the long-term evolution of a gene drive against *UL26* and *UL35*, fibroblasts were co-infected with Towne-GFP and either GD-UL26 or GD-UL35 in a 99/1% ratio, and compared to cells infected with only Towne-GFP (MOI=1). Viral titers and the population of unconverted GFP-only viruses representing unmodified or drive-resistant viruses were followed over time in four or eight independent replicates (Figures 4C-E). In both situations and similarly to previous experiments, the drive first spread efficiently and the proportion of GFP-only viruses reached a minimum of around 20% (for GD-UL35) or 50% (for GD-UL26). In a second phase that corresponded to the positive selection of drive-resistant viruses, the population of GFP-only viruses rebounded and gene drive viruses ultimately represented only a small fraction of the population. Notably, viral titers dropped importantly when the drive reached its maximum, around 100-fold in both cases. This finding confirmed our previous observation that a gene drive with moderate or severe replicative defects could spread in the viral population, and that it could cause an important, albeit transient, reduction of viral titers. Viral levels then increased concomitantly to the rebound of the GFP-only population, but remained two-to five-fold lower until the end of the experiment when compared to cells infected with only Towne-GFP (dashed line in Figures 4D, E). As this remaining population likely represented drive-resistant viruses with a mutated target site, this observation suggested that drive-resistant viruses might have acquired a permanent replicative defect.

### Drive-resistant viruses are durably attenuated

To investigate the population of drive-resistant viruses, viral clones resistant to either GD-UL26 or GD-UL35 and originating from three independent co-infection experiments were isolated and purified. PCR and Sanger sequencing of 11 viral clones for each condition revealed that the target site was mutated as expected (Figures 5A-B). For *UL26*, six clones had an out-of-frame mutation that would disrupt translation, and the five others had an in-frame mutation that nonetheless mutated the conserved proline. On the other hand, every clone resistant to GD-UL35 had a mutated start codon that would prevent translation of the long UL35 isoform. Interestingly, nine of 11 *UL35*-resistant clones had an identical 26 bp deletion that can probably be explained by the presence of microhomology segments on both sides of the cleavage site (Figure 5B). The viral titer of these drive-resistant viruses was then compared to Towne-GFP. The 11 viral clones resistant to GD-UL26 were severely impaired and had a significant 10-fold reduction of viral titers on average, after infection at MOI=0.1 (p=0.0003, Welsh t-test on log-transformed data, Figure 5C). To characterize in detail this replication defect, we selected three resistant clones with in-frame (clones 5, 9, 10) or out-of-frame (clones 3, 7, 11) mutations. Both in-frame and out-of-frame mutants had significantly reduced plaque sizes (p<0.0001, Kruskal-Wallis test, Figure 5D). Besides, multistep growth curves after infection at MOI=0.1 indicated that both in-frame and out-of-frame mutants replicated with a slower dynamic (Figure 5E). Clones with out-of-frame mutations were severely impaired, with a 700-fold reduction at day 3 and 45-fold at day 10 (p<0.0001 for the three time-points, repeated measure two-way ANOVA). In-frame mutants had a more modest defect. Viral levels were significantly reduced 20- and six-fold after three and six days (p=0.002 and 0.0119, respectively, same test as above) and remained attenuated twofold at day 10 (p= 0.2134). This result indicated that viruses resistant to GD-UL26 were significantly attenuated. Importantly, the significant defect of in-frame mutants provided an experimental confirmation that the targeted proline was important for viral fitness.

**Figure 5:**
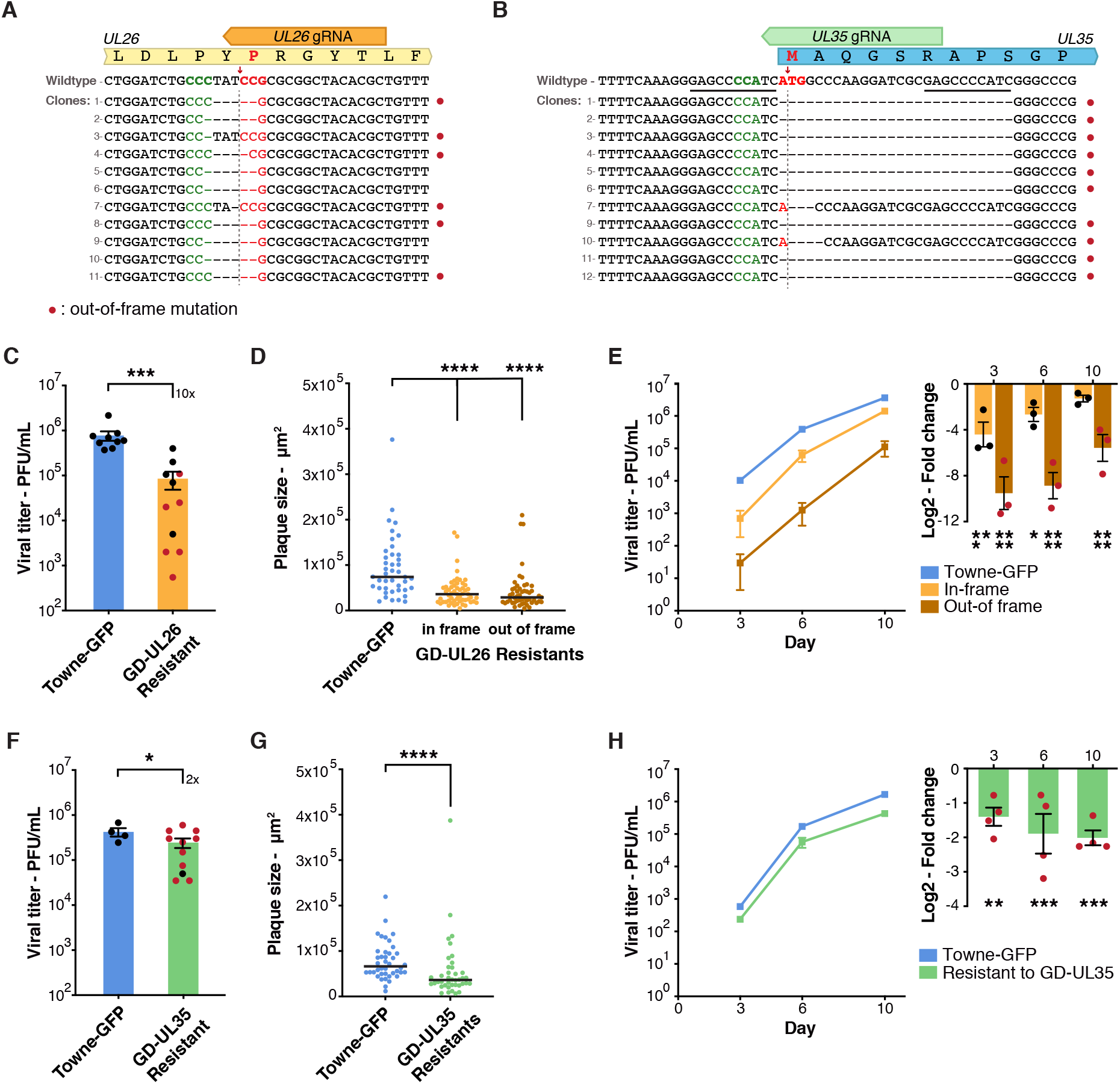
Drive-resistant viruses are significantly attenuated. **A-B.** Sanger sequencing of the target site of 11 viral clones resistant to either GD-UL26 (left) or GD-UL35 (right). CRISPR cleavage sites are shown by red arrows, PAMs are highlighted in green and targeted codons in red. Microhomology sequences around *UL35* cleavage site are underlined in black. **C**. Viral titer in fibroblasts infected with Towne-GFP or viruses resistant to GD-UL26 after 10 days. MOI=0.1, n=9 (Towne-GFP) or n=11 (GD-UL26 resistant). **D**. Plaque size in fibroblasts infected with Towne-GFP or viruses resistant to GD-UL26, after 7 days. Black lines indicate the median, n=40-55. **E.** Multi-step growth curve after infection at MOI=0.1 (left) with the corresponding log2-fold change between mutants and Towne-GFP (right). n=3. **F.** Viral titer in fibroblasts infected with Towne-GFP or viruses resistant to GD-UL35 after 7 days. MOI=1, n=4 (Towne-GFP) or n=11 (GD-UL35 resistant). **G.** Plaque size in fibroblasts infected with Towne-GFP or viruses resistant to GD-UL35, after 7 days. Black lines indicate the median, n=40. **H.** Multi-step growth curve after infection at MOI=0.1 (left) with the corresponding log2-fold change between mutants and Towne-GFP (right). n=4. Replicates with out-of-frames mutations are highlighted in red. Titers were measured by plaque assay and are expressed in PFU per mL of supernatant. Data shows mean and SEM between biological replicates. Asterisks summarize the results of statistical tests. *: p < 0.05, **: p < 0.01, ***: p < 0.001, ****: p < 0.001. (C,F: Welsh t-test on log-transformed data, D,G: Mann-Whitney or Krustal-Wallis test, E,H: two-way repeated-measure ANOVA with Holm-Šidák’s multiple comparisons test on log-transformed data).

Similarly, the 11 viral clones resistant to GD-UL35 had a significant two-fold replicative defect, compared to Towne-GFP after infection at MOI=1. (p=0.0366, Welsh t-test on log-transformed data. Figure 5F). Four resistant mutants (clones 1, 3, 4, 5) with the same 26 pb deletion were selected for further analysis. These drive-resistant viruses had significantly reduced plaque sizes (p<0.0001, Mann-Whitney test, Figure 5G), and replicated with a significant two-to four-fold defect after infection at MOI=0.1 (p=0.0067, 0.008, 0.008 at day three, six and 10, respectively, repeated measure two-way ANOVA. Figure 5H). Altogether, these results indicated that, for both GD-UL26 and GD-UL35, drive-resistant viruses were attenuated compared to Towne-GFP and had acquired long-term replicative defects.

Our results show that a strategy of designing gene drives against conserved and critical viral sequences can counterbalance the evolution of resistance. First, spread of the gene drive in the unmodified viral population caused a strong and transient reduction of viral levels. In a second phase, and even though engineered viruses didn’t stably persist in the viral population, remaining viruses were mutated and had acquired replicative defects, leading to a long-term reduction of viral levels.

## Discussion

In this report, we analyzed the evolution of resistance in a viral gene drive against hCMV. Using multiple examples, we showed that, after the successful invasion of the wildtype population, drive-resistant viruses with a mutated target site are positively selected and outcompete gene drive viruses. These numerical and experimental results mimic perfectly what is observed in insect experiments (14–16): the rapid evolution of resistance prevents the fixation of the modified sequence in the population.

Gene drives in mosquitoes generally follow one of two basic strategies: population-suppression drives aim to eliminate the targeted insect population, whereas population-replacement drives aim to replace wild populations with engineered, often pathogen-resistant, animals (38). Population-replacement drives could be imagined in viruses, where for example a drug-responsive gene would be introduced in the viral population, but here we attempted to design population-suppression viral drives that lead to a significant reduction of viral levels. The difficulty lies in designing gene drive viruses with replicative defects that spread efficiently. Our previous study focused on a gene drive against *UL23*, a viral gene that is dispensable in normal cell culture conditions (6, 11). UL23 is involved in immunity evasion, and the replication of *UL23* mutant viruses is severely impaired by interferon γ. We showed that a gene drive against *UL23* could still spread in the presence of interferon γ, which gave us the first indication that a gene drive with a replicative defect could spread. Here, we designed and tested gene drives against eight additional viral genes that are necessary for efficient viral infection independently of the culture condition (Table 1). We showed that gene drives against the tegument genes *UL26* or *UL35* could spread efficiently in the viral population. GD-UL35 only had a moderate replicative defect, but the GD-UL26 virus was almost non-infectious. However, co-infection with unmodified Towne-GFP virus efficiently complemented the defective viruses and enabled their replication. The modification spread efficiently into the viral population as long as wildtype viruses were present. Once the target site had been mutated, it became unable to propagate further. This represents an important example for the design of suppression drives against herpesviruses.

In insects, targeting evolutionarily conserved sequences, such as the *Doublesex* gene, mitigates the appearance of drive-resistant sequences (19, 39, 40), and our approach followed similar principles. Some regions of the hCMV genome are highly variable, but others show a high degree of sequence conservation, and both *UL26* and *UL35* are reported to be among the most conserved hCMV genes (41). We designed gRNAs in regions of low genetic variation, and targeted sequences encoding conserved amino acids. GD-UL26 and GD-UL35 are examples of gene drives with high and low fitness costs, respectively. As predicted by numerical simulations (Figures 1B, 4D-E), drive-resistant viruses were selected rapidly in the gene drive against *UL26*, and GD-UL26 didn’t reach more than 50% of the population. By contrast, the drive against *UL35* had a higher penetrance and reached up to 80% of the population. In both cases, drive-resistant viruses had mutations that can be predicted to seriously affect the fitness of the virus. Viruses resistant to GD-UL26 had a significant growth defect compared to wildtype viruses, with both in-frame and out-of-frame mutants being attenuated (Figures 5). Absence of *UL35* is reported to cause a modest growth defect (26), and GD-UL35-resistant viruses lacking *UL35* start codon were slightly but significantly attenuated. Mutation of *UL35* had only a moderate effect in our cell culture experiments, but would likely have important consequences *in vivo*. In summary, this work presents important proofs of principle for the design of viral gene drives. It demonstrates that both gene drive and drive-resistant viruses can have replication defects and be durably attenuated, leading to a long-term reduction of viral levels.

We aim to ultimately design similar gene drive systems that could be used to treat herpesvirus diseases. Patients infected with a herpesvirus and unable to control it could be superinfected with a gene drive virus that would reduce the infection. hCMV reactivation in immunosuppressed patients after organ transplant could be one use (42, 43). How a viral gene drive spreads *in vivo* and how the immune system reacts to superinfection will have to be studied in animal models. Our work nonetheless brings important considerations about viral dynamics. In particular, in our gene drives against *UL26* or *UL35*, viral levels dropped importantly when drive penetrance reached its maximum, with a 10–100-fold reduction of viral titers (Figures 4D-E). This transient drop of viral levels could have important implications *in vivo*, as it could give the immune system a transient window to control the infection. Similarly, even a small decrease of viral fitness caused by drive-resistant mutations could have huge benefits for the infected patient. Indeed, *in vivo*, a successful gene drive wouldn’t require reducing viral levels significantly by itself, but to do so just enough for the immune system to take control of the infection.

Pre-existing genetic variation in hCMV or other herpesviruses would hamper the capacity of a gene drive to efficiently target wild viruses in infected patients. We designed CRISPR gRNAs in regions of low genetic diversity, but the number of available genomes (235) is small compared to the size of the hCMV population. A better assessment of the genetic diversity of herpesviruses will be necessary before a gene drive can be used against wild viruses. Nonetheless, one can envision that patients could be treated with an array of gene drive viruses, each targeting different variants or different locations. Such a strategy would increase redundancy and limit the probability that variants could escape the drive.

As a final note, the development of gene drives raises important ecological and biosafety concerns, and our approach follows the guidelines established by the NIH and the National Academy of Science (44, 45). In particular, our work was conducted using laboratory viral strains unable to infect human hosts (46), and thus, eliminated risks of inadvertent release of gene drive viruses into the wild.

## Material and Methods

### Cells and viruses

Human foreskin fibroblast cells were obtained from the ATTC (#SCRC-1041) and cultured in DMEM (10-013-CV, Corning, Corning, NY, USA), supplemented with 10% FBS (Sigma-Aldrich, USA) and 100 μm/L penicillin/streptomycin (Corning, USA). Cells were regularly tested negative for mycoplasma and used between passages 3 and 15.

hCMV TB40/E-Bac4 (47) and Towne-GFP (T-BACwt)(20) were kindly provided by Edward Mocarski (Emory University, USA). Viral stocks were prepared and plaque assays performed exactly as reported (6).

Co-infection experiments were performed by co-infecting confluent fibroblasts with wildtype Towne-GFP and gene drive viruses for 1 h, with a total MOI of 0.1–1. Experiments were conducted using 12-well plates with 1 mL of medium per well. For time-course experiments over multiple weeks, 100–200 μL of supernatant was used to inoculate fresh cells for 1 h before changing media, maintaining an MOI around 1 at each passage. Viral titers were measured at each passage.

For plaque size analysis, images of fluorescent viral plaques were acquired with a Nikon Eclipse Ti2 inverted microscope and Nikon acquisition software (NIS-Element AR 3.0). Plaque size in pixels was measured using ImageJ (v2.1.0) and then converted to μm^2^.

### Cloning and generation of gene drive viruses

The gene drive construct against *UL23* was as described (6). The core gene drive cassette comprises a codon-optimized *SpCas9* (from *Streptococcus pyogenes*), followed by an mCherry fluorescent reporter and a U6-driven gRNA (Figure 2A). This cassette is surrounded by homology arms specifics to the site of integration. Gene drive plasmids against *UL122, UL79, UL99, UL55, UL26, UL35, UL82* and *UL69* were built by serial modifications of the *UL23* gene drive plasmid. Briefly, homology arms and the gRNA for *UL23* locus were removed by restriction enzyme digestion, and replaced by new homology arms and gRNAs by Gibson cloning (NEB, USA), using PCR products or synthesized DNA fragments (GeneArt™ String™ fragments, ThermoFisher, USA). GD^Towne^-UL23 was generated with a gene drive plasmid targeting *UL23* CDS.

Cell lines stably expressing *UL26* or other viral genes were generated using lentiviral constructs. Lentivirus expression plasmids were cloned by serial modifications of Addgene plasmid #84832 (48), using digestion/ligation and Gibson cloning. The final constructs expressed the viral gene of interest, followed by in-frame puromycin and BFP reporters interleaved with self-cleavable 2A peptides under an EF1a promoter. Lentiviruses were produced in HEK293 cells using standard protocols as reported (49). Fibroblasts were then transduced with lentiviruses and selected with 1 μg/mL puromycin for 1 week before being used (ant-pr-1, Invivogen, USA).

Purification of gene drive viruses was performed as reported (6). Briefly, fibroblasts were transfected by nucleofection (Kit V4XP-2024, Lonza, Switzerland) with the gene drive plasmid, and infected 48 h later with Towne-GFP virus. Recombinant viruses expressing mCherry were isolated and purified by several rounds of serial dilutions and plaque purification.

Viral clones resistant to either *UL26* or *UL35* gene drives were isolated by plaque purification of GFP-only viruses at the end of co-infection experiments. Mini-stocks were titrated by plaque assays, and the sequence of the target site was analyzed by PCR and Sanger sequencing. (UL26 primers: F: GGCGCGTTATAAGCACCGTGG, R: GCCGATGACGCGCAACTGA; UL35 primers: F: ACGTCACTGGAGAACAATAAAGCGT, R: GGCACGCCAAAGTTGAGCAG). 12 resistant clones originating from 3 independent experiments were first isolated, but in both cases, only 11 could be successfully purified and sequenced.

### Amplicon sequencing

Total DNA was extracted from infected cells with Qiagen DNeasy kit. A 470 bp PCR product surrounding the *UL23* cut site was amplified using Phusion high-fidelity polymerase (NEB, USA) and column purified (Macherey-Nagel, Germany). Primers contained a 5’-overhang compatible with Illumina NGS library preparation. Amplicons were pooled and sequenced on an Illumina Miseq (2×300 paired-end). Library preparation and sequencing were performed by SeqMatic (Fremont, CA, USA). Analysis of genome editing outcomes from sequencing data was generated using CRISPResso2 software pipeline (34). Forward primer: TCGTCGGCAGCGTCAGATGTGTATAAGAGACAGGCTTGGGGCATAAAACACCG; Reverse primer: GTCTCGTGGGCTCGGAGATGTGTATAAGAGACAGCCCAGGTACAGTTCAGACGG.

### Sequence alignment and motif analysis

Protein sequences of the herpesviridae US22 protein family were downloaded from Uniprot and curated to remove duplicates. Motif discovery among the 31 protein sequences was performed using the Meme Suite 5.3.0 (http://meme-suite.org/), using OOPS parameter and looking for motifs 10–30 in lengths (50).

Full-length genome sequences of 235 hCMV clinical and laboratory strains were downloaded from the NIAID Virus Pathogen Database and Analysis Resource (ViPR: https://www.viprbrc.org/)(51). The frequencies of variants around the CRISPR target site were calculated using simple Python and R scripts. Briefly, we recovered the sequences around the target site, using BLAST locally with the sequence of the target site as the query and the 235 hCMV sequence as the search database. Base frequencies on the aligned sequences were calculated and plotted using R packages APE v5.3 and ggplot2 v3.3.0 (52, 53).

### Statistics and reproducibility

Plaque assay data do not satisfy the normality condition required for parametric tests, and F-tests further showed that variances were often non-homogeneous. As a consequence, statistical tests on plaque assay data were performed on log-transformed data, using statistical tests that didn’t assume homogeneity of the variance. Namely, we used Welsh t-test to compare two groups, and Welch and Brown-Forsythe ANOVA followed by Dunnet’s T3 comparisons test to compare 3 or more groups. To compare multi-step viral growth curves in figure 5, we used repeated-measure two-way repeated-measure ANOVA followed by Holm-Šidák’s multiple comparisons test. Differences in viral plaque size were analyzed using Mann-Whitney or Kruskal-Wallis nonparametric tests. Analyses were run using GraphPad Prism version 8.1.1 for macOS (GraphPad Software, USA, www.graphpad.com). Exact p-values and summaries are reported in the text and the source data, respectively.

### Numerical simulations

Numerical simulations of viral gene drive were computed using a simplified viral replication model. Briefly, in each viral generation, N virtual cells were randomly infected and coinfected by N viruses (MOI =1), producing a new generation of viruses. In this new generation, wildtype viruses co-infected with drive viruses were converted to new gene-drive viruses or resistant viruses with a ratio of 90/10. Gene drive viruses replicate with a fitness cost f, and the coinfection rate is calculated from the MOI assuming a Poisson distribution. The code and a more thorough description are available at https://github.com/mariuswalter/ViralDrive.

## Data and code availability

The data supporting the findings of this study are available within the paper and its supplementary files. Amplicon Sequencing data have been deposited in the Short Read Archive with BioProject accession no. <u>PRJNA556897</u>. Lentivirus expression plasmids for *UL23, UL122, UL79, UL99, UL55 and UL26* will be deposited in Addgene. Viruses and other reagents developed in this study are available upon request and subject to standard material transfer agreements with the Buck Institute. Source data are provided with this manuscript. Any other relevant data are available upon reasonable request. Code developed for numerical simulations is available on GitHub (https://github.com/mariuswalter/ViralDrive).

## Acknowledgments

We thank members of the Verdin laboratory for technical and conceptual help. This study was funded through institutional support from the Buck Institute for Research on Aging.

## Author contributions

M.W. designed the study. M.W. and R.P. conducted experiments. E.V. supervised and funded the project. M.W. and E.V. wrote the manuscript.

---

**Supplementary Figure S1:**
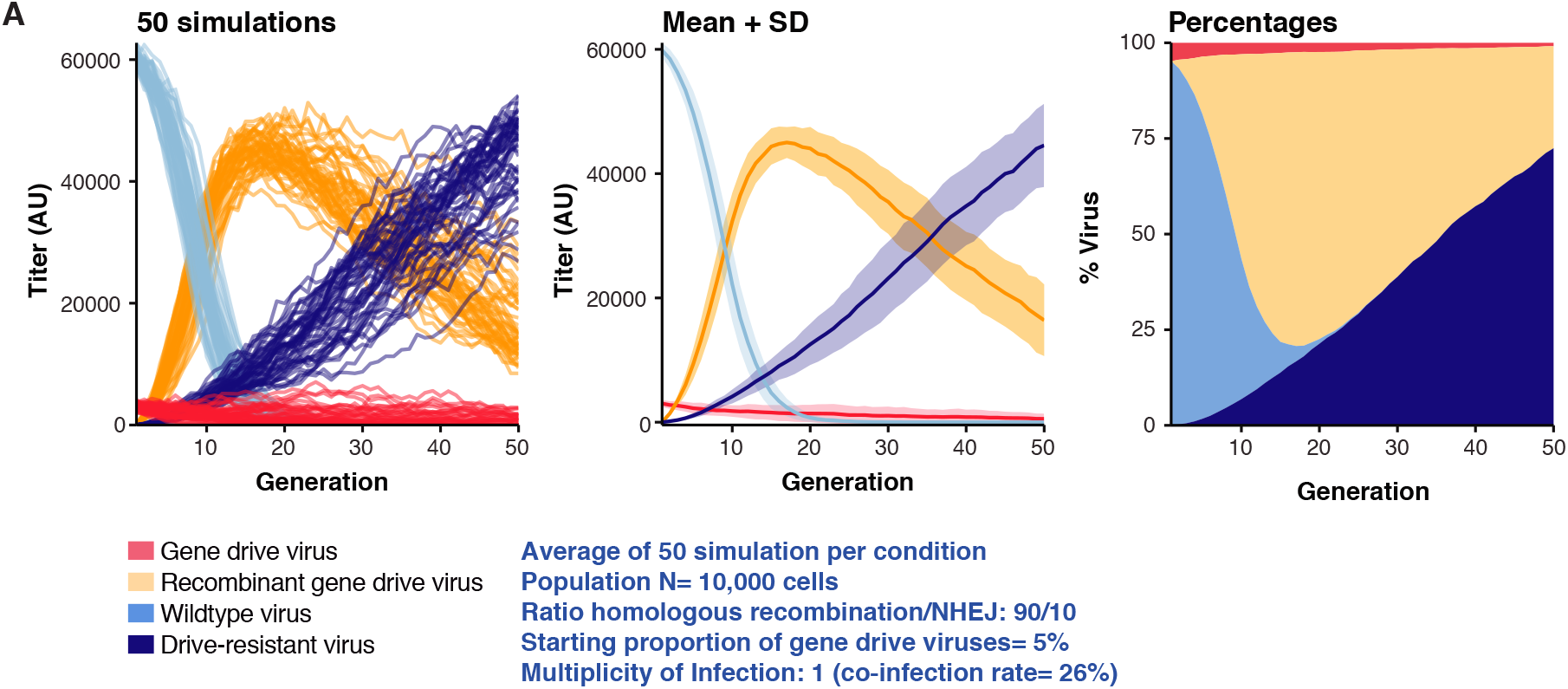
Example of numerical simulation. Example of a simulation with a replicative fitness f=0.9 for gene-drive viruses. Left panel shows 50 independent simulations. Middle: mean and standard deviation of the 50 simulations. Right: mean proportion of the different viruses.

**Supplementary Figure S2:**
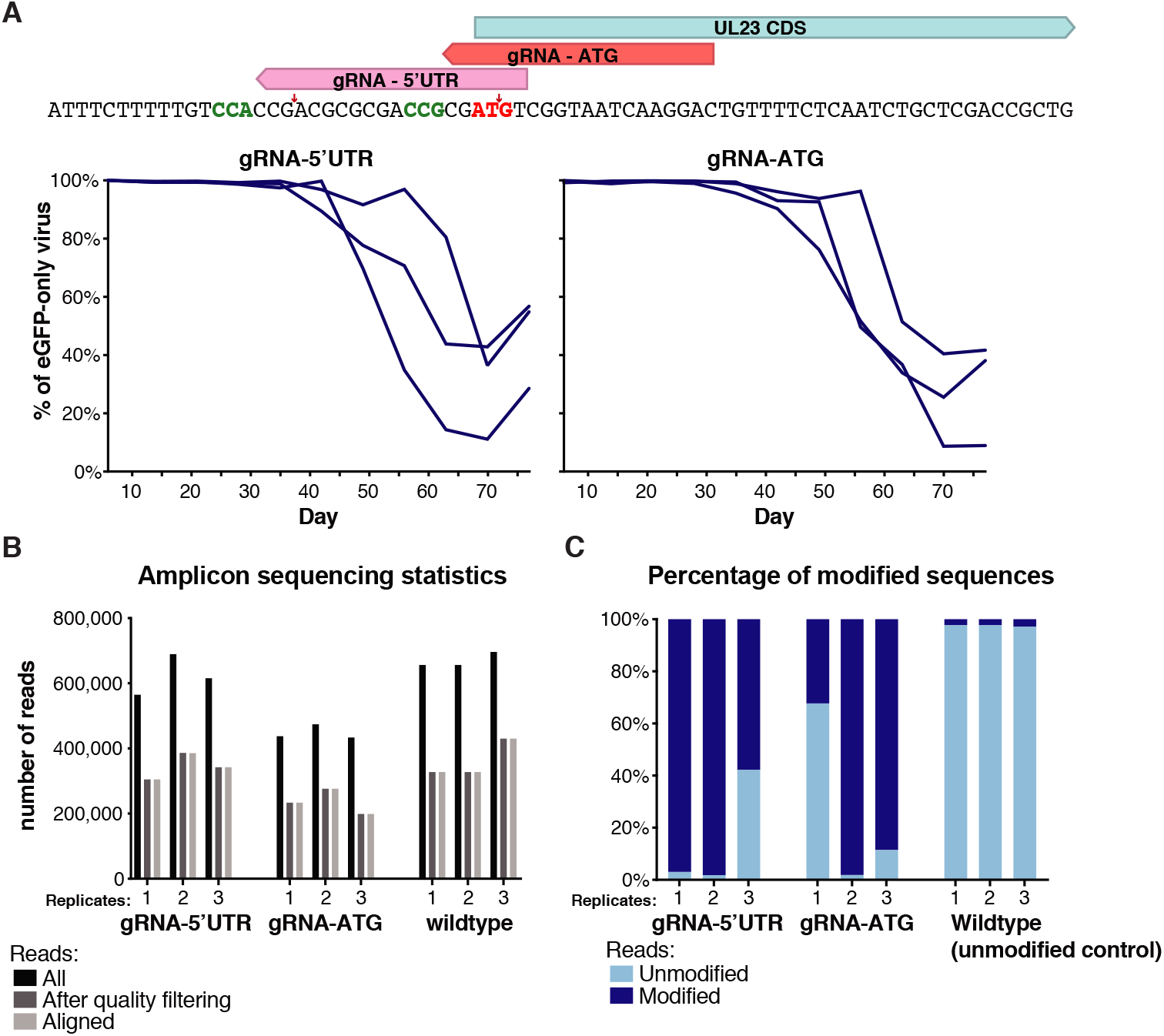
Amplicon sequencing of drive-resistant target sites. **A**. Proportion over time of viruses expressing only GFP in a gene drive against *UL23* 5’UTR or ATG. Fibroblasts were initially transfected with a gene drive plasmid against UL23 5’UTR or UL23 start codon, and subsequently infected with Towne-GFP. **B**. Amplicon-sequencing statistics of three biological replicates, at day 70. Towne-GFP-infected cells were also sequenced as an unmodified control. **C**. Proportion of edited genomes after amplicon sequencing. Same as figure 2E, but including the wildtype control.

**Supplementary Figure S3:**
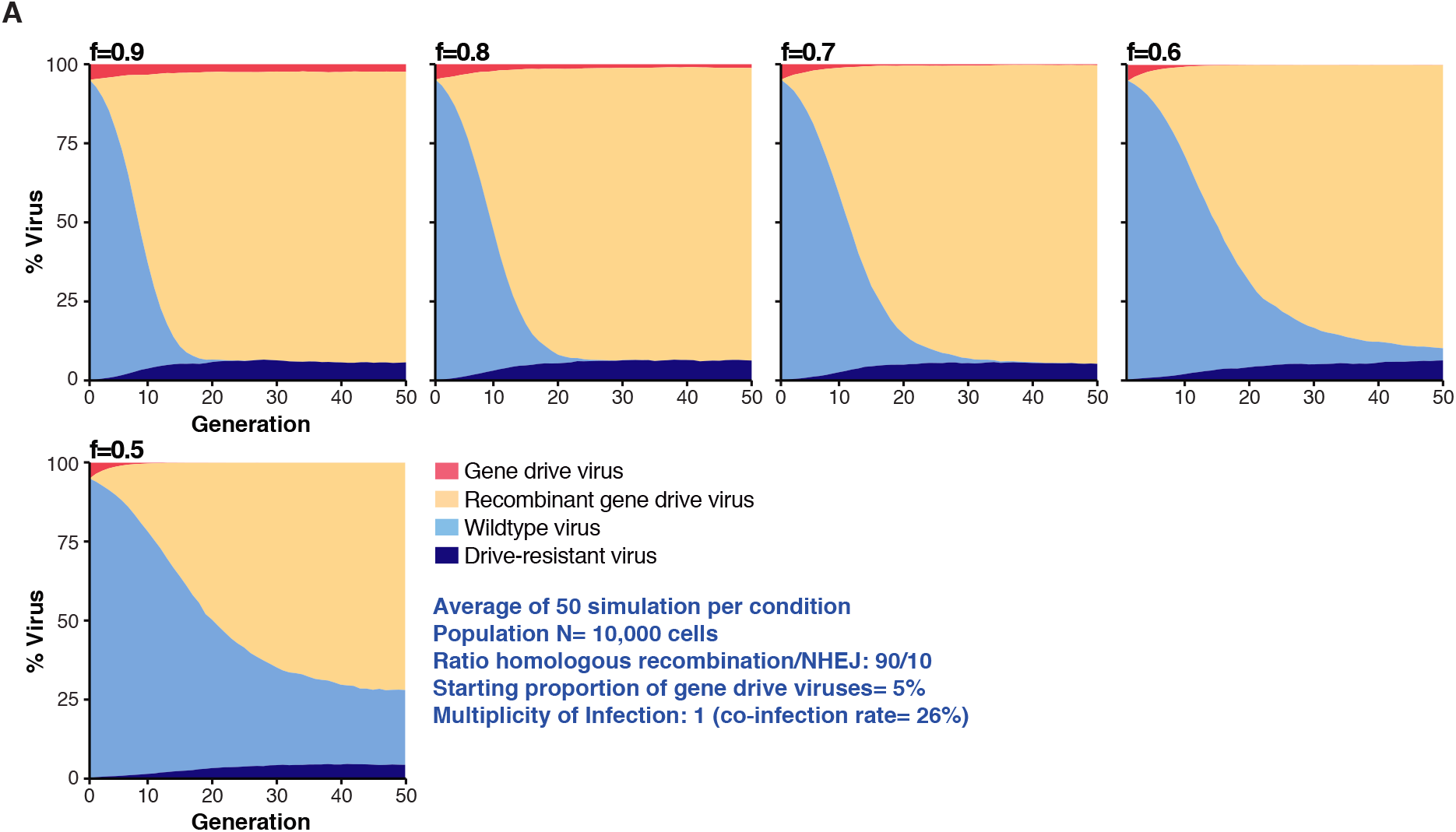
Numerical simulations with defective drive-resistant viruses. Numerical simulation with gene-drive and drive-resistant viruses having the same fitness f. In these simulations, at each viral generation, N virtual cells were randomly infected and coinfected by N viruses, producing a new generation of viruses. When a cell was coinfected by wildtype and gene-drive viruses, wildtype viruses are converted to new gene drive viruses or resistant viruses in a 90/10 ratio.

**Supplementary Figure S4:**
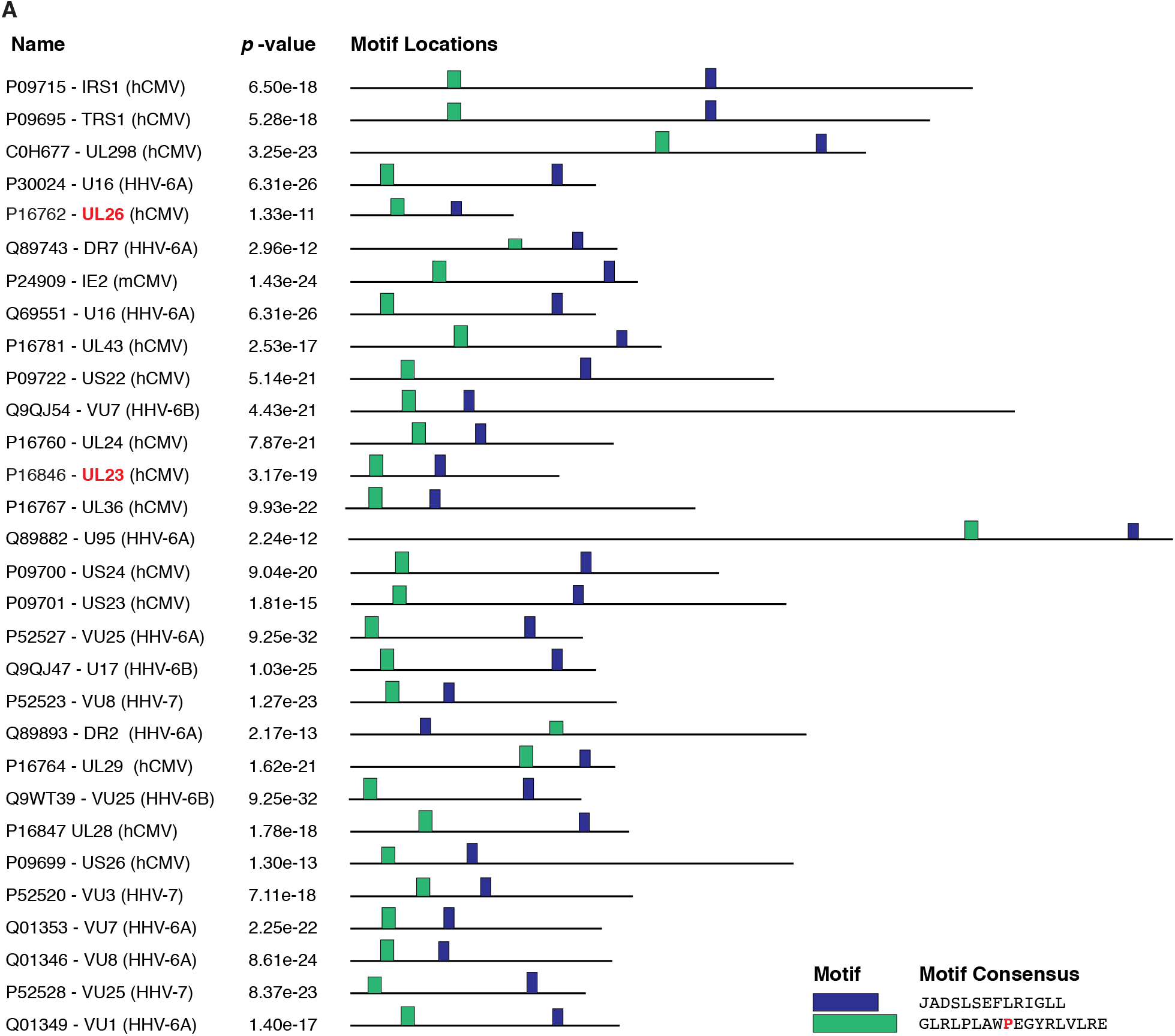
Localization of conserved motifs in the US22 protein family. Localization of conserved motifs in 31 proteins of the US22 family of herpesviral proteins. First column gives Uniprot IDs, gene names and viruses of origin.

